# Short-term Hebbian learning can implement transformer-like attention

**DOI:** 10.1101/2023.05.31.543109

**Authors:** Ian T. Ellwood

**Affiliations:** Department of Neurobiology and Behavior, Cornell University, Ithaca, NY, 14853, USA

**Keywords:** Transformers, Attention, Hebbian Learning, NMDA Receptors, Short-Term Synaptic Plasticity

## Abstract

Transformers have revolutionized machine learning models of language and vision, but their connection with neuroscience remains tenuous. Built from attention layers, they require a mass comparison of queries and keys that is difficult to perform using traditional neural circuits. Here, we show that neurons can implement attention-like computations using short-term, Hebbian synaptic potentiation. We call our mechanism the match-and-control principle and it proposes that when activity in an axon is synchronous, or matched, with the somatic activity of a neuron that it synapses onto, the synapse can be briefly strongly potentiated, allowing the axon to take over, or control, the activity of the downstream neuron for a short time. In our scheme, the keys and queries are represented as spike trains and comparisons between the two are performed in individual spines allowing for hundreds of key comparisons per query and roughly as many keys and queries as there are neurons in the network.

## Introduction

The transformer architecture (1) has demonstrated remarkable, seemingly cognitive-like abilities, including few-shot and single-shot learning, puzzle solving, computer programming, image generation and style transformations (2–8). While still falling short of human abilities on many tasks, transformer-based models continue to improve, and it is natural to ask if their architecture may have relevance to the computations performed by neural circuits in the brain.

The new ingredient in transformers that differs from alternative deep learning methods like recurrent neural networks and convolutional neural networks is the attention layer (1, 9). This layer acts on three collections of vectors known as queries, keys and values. Each query is compared with each key and the output associated with each query is a weighted sum of the values associated with the best matching keys (Figure 1 A, B). In state-of-the-art transformers, the number of queries and keys can be in the tens of thousands.

**Fig. 1.**
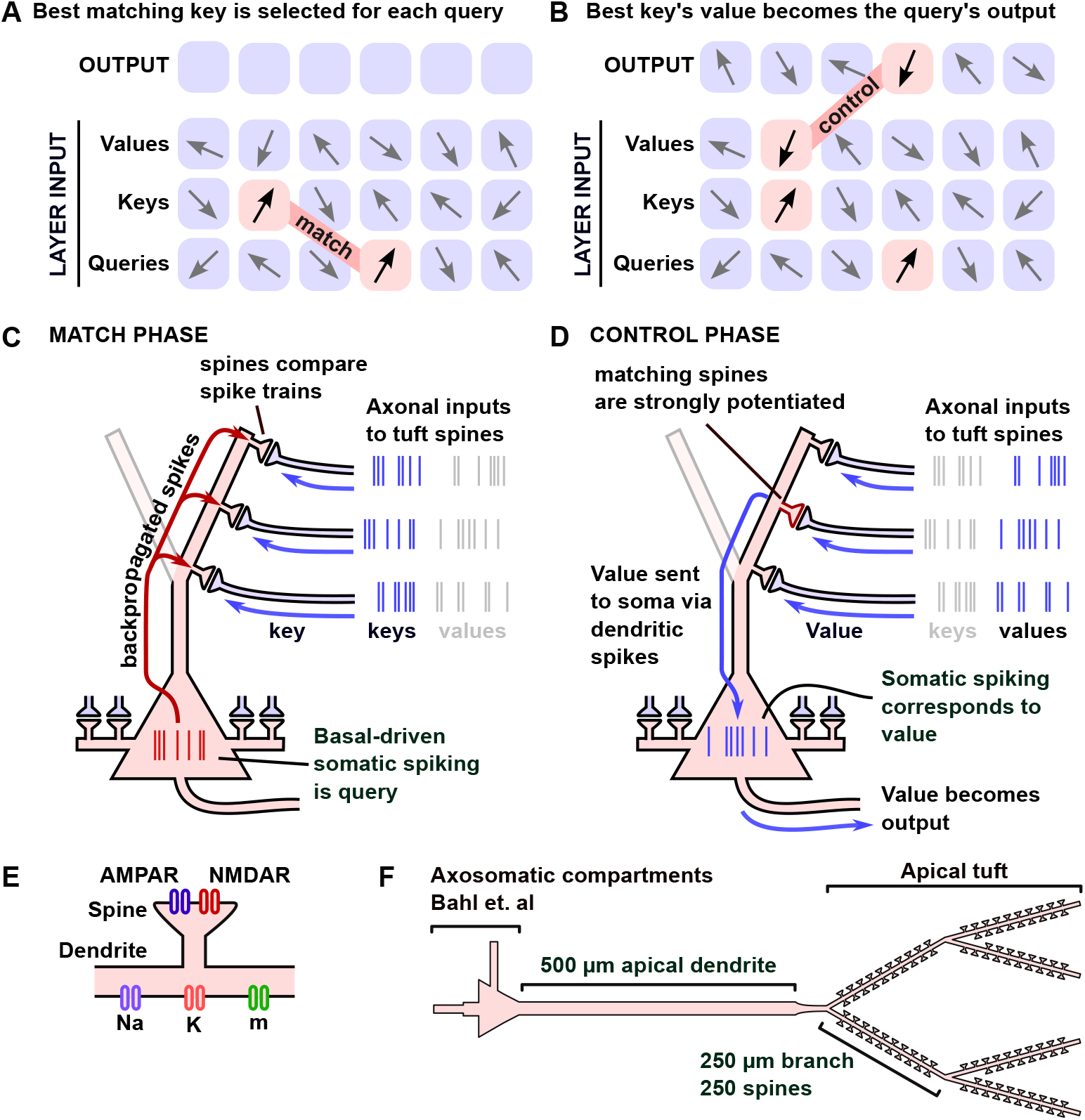
Overview of the match-and-control principle. **A)** In an attention layer, each query is matched with the nearest key. **B)** The output associated with the query is given by the value associated with the closest matching key. **C)** The Match Phase: We propose that somatic activity, driven by basal dendrites, represents a query. This spike train backpropagates into the apical tuft, where it is compared in dendritic spines with axonal activity, representing keys. **D)** The Control Phase: Spines that received matching somatic and axonal activity potentiate. Subsequent axonal activity at potentiated spines represents the key-associated value and is transmitted to the soma via dendritic spiking. **E)** The ion channels in the apical tuft and spines of the pyramidal neuron model. **F)** The dimensions of the full pyramidal neuron model.

This matching of multiple queries and keys has no known analogue in the brain, but a related idea is the comparison between a single query and many keys. This simpler computation is similar to models of memory recall, where a query is analogous to a contextual cue and is used to find the best stored key or memory in an associative neural network, such as a modern Hopfield network (10, 11). This analogy has been explored thoroughly in (12), while (13) made the connection with hippocampal circuits more explicit. (See also (14, 15) for two very different proposals and (16) for efforts to find transformer-like representations in the brain.)

We note, however, two important differences between these models and transformers. First, only a single query is compared with the collection of keys. While one can imagine sequentially comparing queries or that multiple regions of the brain can receive different queries simultaneously, in either case, the number of queries will be far smaller than the number of keys. In contrast, in transformers, queries and keys are typically of the same quantity. A second issue is that the keys in these models are not dynamic, but fixed vectors learned through experience, as they represent stored memories in the hippocampus. While new keys can be added or modified through learning, this largely fixed collection of keys is a departure from the spirit of the transformer where the keys, queries and values are generated on the fly with each pass through the network.

In the hippocampal-inspired models of transformers, the vector-valued queries are represented as a collection of firing rates in a bundle of axons, as is typical for machine-learning-inspired models of the brain. However, using firing-rate vectors runs into difficulties when many queries and keys must be compared simultaneously. For example, suppose one built a model in which a bundle of axons, representing a key, is sent to multiple brain regions so that it may be compared with multiple queries. Even though the activity in each copy of the bundle of axons is identical, each brain region will receive a different message, transformed by the synaptic weight matrix between the axons and the downstream neurons. To implement an attention-like computation, these weight matrices must be nearly identical, or each comparison will be made with a transformed key. Fixing this issue would require sophisticated learning rules that are unknown in the brain.

An alternative, which we examine in this study, is that queries and keys are represented by short trains of spikes, as has been explored in recent attempts to implement transformers on neuromorphic chips (17–19). An advantage of this choice is that every neural receiver of a train of spikes will receive the same message, regardless of the synaptic strength of the connection (provided that the strength is not zero). In addition, if one wishes to send a vector to many receivers in the brain, one can use the natural arborization of axons without any additional machinery.

A final advantage of using spike trains is that biological mechanisms that compare two spike trains are well-established in neuroscience. When a neuron receives a train of spikes from an excitatory axon, glutamate is released from the axon terminal and binds with glutamate receptors on the postsynaptic spine. Of particular interest, NMDA receptors only open and allow calcium entry into the spine when they are bound to glutamate and the spine is depolarized, which can occur when somatic action potentials backpropagate into the dendritic tree (20). Under the right conditions, the amount of calcium entry into a spine can thus act as a proxy for how similar the pattern of presynaptic action potentials is to the pattern of somatic spikes.

In standard theories of synaptic plasticity, when large amounts of calcium enter the spine, the spine will undergo long term potentiation (LTP), strengthening the synapse (21, 22). This mechanism is believed to be the biological basis for Hebbian learning in the brain. Here, we consider a modification of these rules in which the potentiation following calcium entry is larger than what is typically found in LTP experiments, but is transient, lasting only seconds, making it a form of short-term plasticity. If this potentiation is large enough to allow a single axon to initiate dendritic spikes, we show that a computation similar to transformer attention can be performed. We emphasize that because the changes to synaptic weights are not long lasting, our scheme is an example of synaptic computation (23), not learning or memory.

## Results

### The Match-and-Control Principle

In attention layers, when a key matches with a query (i.e., the vectors have a large dot-product), the value associated with the key becomes the output associated with the query. We emulate this process using pyramidal neurons as follows. We suppose that a train of spikes in the soma of a neuron represents a query vector. These somatic spike trains are generated by axonal inputs to the basal dendrites of the neuron. We represent multiple queries with multiple neurons. The key vectors are represented by trains of action potentials carried by individual axons (one axon per key) that synapse onto the apical dendritic tree of the neuron.

During the match phase (Figure 1C), the firing patterns in the axons synapsing onto the apical dendrites are compared with the back-propagated action potentials (BAPs) evoked by basal-driven activity, using the amount of calcium entry into the apical dendritic spines. In detail, in each spine, NMDA receptors open when depolarizations from the BAPs occur just after glutamate is released onto the post synaptic density of the spine. Since the pattern of glutamate release is determined by the pattern of presynaptic spikes, which we identify with the key, the total amount of calcium that enters the spine acts as a measure of the similarity of the timing of spikes in the query and key.

One subtlety with this discussion is that small fluctuations of the dendritic membrane potential can allow NMDA receptors to leak sizable amounts of calcium into the spine, spoiling this connection. This can be resolved by considering a calcium detector that is a non-linear function of the concentration of calcium, suppressing responses to small amounts of calcium entry. Such non-linearities can occur when the molecule that senses calcium has multiple binding sites for calcium, all of which must be bound for the molecule to be active. For example, one of the primary molecules that detects spine calcium levels is calmodulin (CaM), which requires four calcium ions to become fully active.

We thus consider a simple polynomial non-linearity and use the fourth power of the calcium concentration as our measure of similarity. Raising the calcium concentration to this power suppresses small fluctuations in calcium and has the nice feature that it narrows the brief calcium entry events when BAPs coincide with glutamate release. We note that in vivo mechanisms for calcium detection are likely more complex than this simple model. For example, CaM’s calcium-binding affinity can be affected by CaM-binding proteins and is not independent for the four calcium binding sites (24, 25). Moreover, there are multiple spine mechanisms for the detection of calcium, including calcium-dependent ion channels that may play a role in this process, each of which can have a different calcium-dependent activity. Nonetheless, all we need for our mechanism to function is any non-linearity that suppresses the response to low levels of calcium entry.

Following the match phase, we propose that spines with high levels of calcium rapidly and transiently potentiate their synaptic strength with their respective axons. In our model, we will remain agnostic about how exactly this potentiation is implemented at the synapse and, as far as we are aware, there is no experimental evidence of rapid large increases in synaptic strength that depend on the postsynaptic potential. In the discussion, we will examine the relative merits of several possible mechanisms, including rapid AMPA phosphorylation, calcium-dependent non-specific ion channels, presynaptic facilitation, post-tetanic potentiation and NMDA spikes. For now, though, we simply assume that some mechanism exists and show its advantages for implementing the attention computation.

In attention layers, a softmax function is used to output a weighted sum of the values that closely match the query. In early attention layers, this process typically returns an average of many values, but in the middle layers, the values are often associated with a single key (12). It is this highly selective form of attention that we wish to emulate, as we feel that it is the most “transformer-like”. Moreover, implementing a softmax across spines is challenging because it requires knowledge of the total amount of potentiation across all the spines to normalize the net output. While this is potentially possible using a more sophisticated setup, here we simply pick a high threshold for integrated calcium entry so that only a very small number of spines ever become potentiated. While this implies that sometimes we will fail to have any match, such failures are a familiar situation in neuroscience. For example, an attempt to recall a memory from a cue can result in no memory being recalled or an attempt to initiate an action can fail to initiate any action.

Once any spines that cross our threshold are potentiated, we enter the control phase (Figure 1D) in which axons presynaptic to potentiated synapses are able to drive spiking in the soma of the postsynaptic neuron, thus taking control of its activity. The same pre-dendritic axons that previously transmitted the keys now transmit spike trains representing key-associated values. The keys and values are thus temporally concatenated in the train of spikes delivered by the axons. To transmit the values associated with the best matching keys, the axons that synapse onto the potentiated spines elicit dendritic spikes that propagate to the soma of the neuron leading to somatic and then axonal spiking. In this way, the somatic activity of the neuron transitions from being the query in the matching phase to being the value associated with the best matched spines in the control phase.

We note that the number of queries in this scheme is equal to the number of participating neurons, while the number of keys is given by the number of presynaptic axons, allowing for large numbers of queries and keys. However, not every key is compared with every query. Dendritic trees typically have thousands of spines, but each axon can make multiple synapses with a single neuron (26–28), making the number of comparisons as much as an order of magnitude smaller than the number of spines and far smaller than the number of neurons in a typical neural circuit, which can range from the thousands to millions. Our proposal should thus be thought of as a kind of “sparse attention” where the number of keys and queries can be enormous, potentially far larger than modern transformers, but only a fraction of them are compared with each other.

### Biophysical model implementing the Match-and-Control Principle

To test if our scheme is biophysically plausible, we implemented a cable-theory based pyramidal neuron model with Hodgkin-Huxley-style ion channels using the NEURON simulator (29) (Figure 1 E, F). In the construction of our model, we aimed to build as simple a model as possible that could implement the match-and-control principle. Real pyramidal neurons include numerous additional channels not considered here, including calcium channels, calcium-activated non-specific ion channels, multiple species of potassium channels and chloride channels. Our model should thus be considered a proof-of-principle, as well as a demonstration that very few ingredients are necessary to produce our desired effects, but not an attempt to implement any specific pyramidal neuron type in the brain.

For the soma, basal dendrites and axon, we used a reduced model of a layer V pyramidal neuron by Bahl et. al. (30). For the dendrites, we built a 500 µm apical dendrite based on a model by Hay et. al. (31) connected to a branching tuft consisting of six 250 µm segments with 250 spines per branch, adding up to a total of 1500 spines. For AMPA and NMDA channels we used models based on (32–34). We assumed that each of 150 axons synapsed 10 times on the dendritic tree, and that they synapsed on neighboring spines to give them the best ability to depolarize the dendrites (28, 35, 36). Our model thus allows 150 unique keys and associated values per neuron. The amount of redundant axonal input on neighboring spines does not have a large impact on the model’s performance as it simply increases the size of the dendritic EPSPs by a factor of 10 for each presynaptic spike in both the match and control phases. This increase can be achieved in a model where each axon only synapses once onto the dendritic tree by rescaling the AMPA conductance, and it is only the product of the AMPA conductance and axon synapse redundancy that is important for the simulations. Nonetheless, we include this feature as it has been observed in vivo and proposed as a mechanism for dendritic spike initiation by single axons.

We used the same ion channels for the apical shaft and tuft dendrites as Hay et. al. (31), but increased the conductance of the fast sodium channels and voltage dependent potassium channels, as their model does not support propagating dendritic spikes. In a short segment connecting the apical dendrite with the tuft branches, we used a slightly higher density of fast sodium channels to allow the dendritic spikes to propagate into the apical shaft. We also increased the axial resistance in this segment to prevent the larger apical dendrite shaft from shunting any currents generated in nearby spines. We note that in some pyramidal neurons, this region would contain a high density of high voltage activated calcium channels (37), but to keep our model simple, we have not modelled calcium dynamics outside of the spines.

### Calcium entry as a measure of spike train similarity

We began by testing whether calcium can be used as a proxy for spike train similarity in our model, as in models of spike-timing dependent plasticity (32, 38). An example simulation is shown in Figure 2A for a spine whose inputs match the somatic action potentials and for a spine whose inputs are random. As expected, matching the glutamatergic inputs to the spine with the BAPs led to increased calcium entry, but we also observed sizable amounts of calcium entry away from BAPs in the spine with unmatched inputs. As described above, using the fourth power of the calcium concentration suppressed these events (Figure 2A, bottom two traces).

**Fig. 2.**
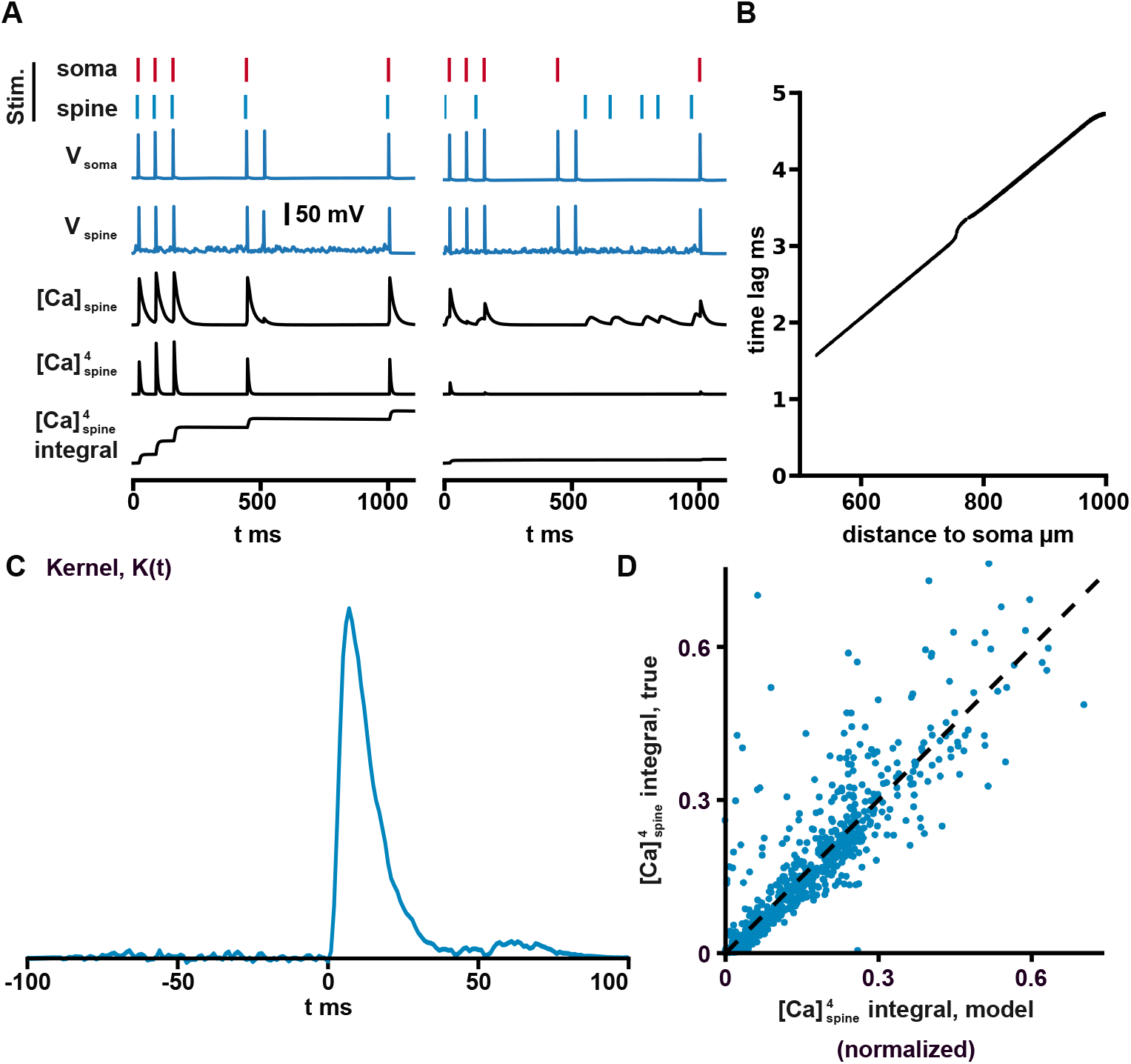
Calcium as a measure of spike train similarity. **A)** An example simulation of the model is shown. The left column of traces are recordings from a spine receiving presynaptic spikes that occurred 7 ms before the back-propagating action potentials arrived from the soma. The right column shows recordings from a spine that received random glutamatergic inputs drawn from an 6 Hz Poisson process. Note that even unmatched glutamatergic inputs produced calcium entry (third trace from bottom, right column), but that raising the calcium concentration to the fourth power suppressed these events (bottom two traces). **B)** The latency for a back-propagated action potential to reach each of the 1500 spines in the dendritic tree (average of 10 simulations). Note that slight differences in time to reach the symmetric branches of the dendritic tree arose in the simulation because the dendrites were receiving random axonal inputs. **C)** Plot of the best-fit kernel used to model the integral of the fourth power of the calcium concentration as a bilinear overlap between the presynaptic and post-synaptic spike trains. Note that the Kernel included a double sigmoidal window that forced its values to exponentially decay to zero outside of the interval [- 30 ms, 30 ms] with a time-constant of 10 ms. **D)** A comparison of the simple kernel model with the simulated integrated calcium signal on a test dataset not included in the fit. 1000/15000 randomly selected test data points are shown.

In attention layers, the similarity of a key and query is measured using a dot product. We thus tested if the net calcium entry could be approximated by a similar bilinear form. To account for the temporal kinetics of NMDA channels and the finite width of BAPs, we used a model with an arbitrary temporal kernel,

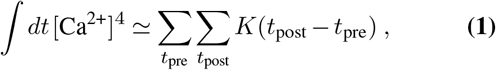

where *t*_pre_ and *t*_post_ are the times of the pre- and postsynaptic BAPs and *K* is the kernel. We note that for *t*_post_ we used the time when the BAP reached the spine, accounting for the transmission delays from the long dendrites of the apical tree shown in Figure 2B.

To estimate *K*(*t*), we used 1 second spike trains sampled from a Poisson process with an average spike rate of 6 Hz and a spike refractory period of 50 ms. We ran 100 simulations, giving a dataset of 15000 presynaptic spike trains and 100 post synaptic spike trains. Using gradient descent on a least squares loss, yielded the best fit for *K*(*t*) shown in Figure 2C. The quality of the fit is shown for a separate test dataset in Figure 2D, where the model explained 81% of the variance.

If the kernel was a delta-function, it would imply that the calcium integral is approximately equal to the ordinary Euclidian inner product between the two spike trains considered as temporal vectors (i.e., vectors with ones where the spikes are and zeros where there are no spikes). Because of the finite width of the peak around zero, the model may be considered a temporally smeared version of a simple dot product.

In this comparison, we removed a small collection of outliers (84/15000 (0.6%) and 77/15000 (0.5%) for training and test data) whose integrated calcium entry was 4 standard deviations larger than the average. Such outliers can occur when the random fluctuations of the membrane potential allow for non-trivial NMDA channel conductance or when spontaneous dendritic spikes occur. Though such events are rare, they are unavoidable in our models due to the large number of random synaptic inputs to the dendrites and reduce the selectivity of our matching procedure.

### Finding the best matched spike train

We measured the optimal time for the presynaptic spikes to arrive by computing the integral of the fourth power of the calcium concentration as a function of the offset between pre- and postsynaptic spikes (Figure 3A). This computation showed that, ideally, presynaptic spikes arrive around 7 ms before the BAPs arrive, a value we use throughout our simulations.

**Fig. 3.**
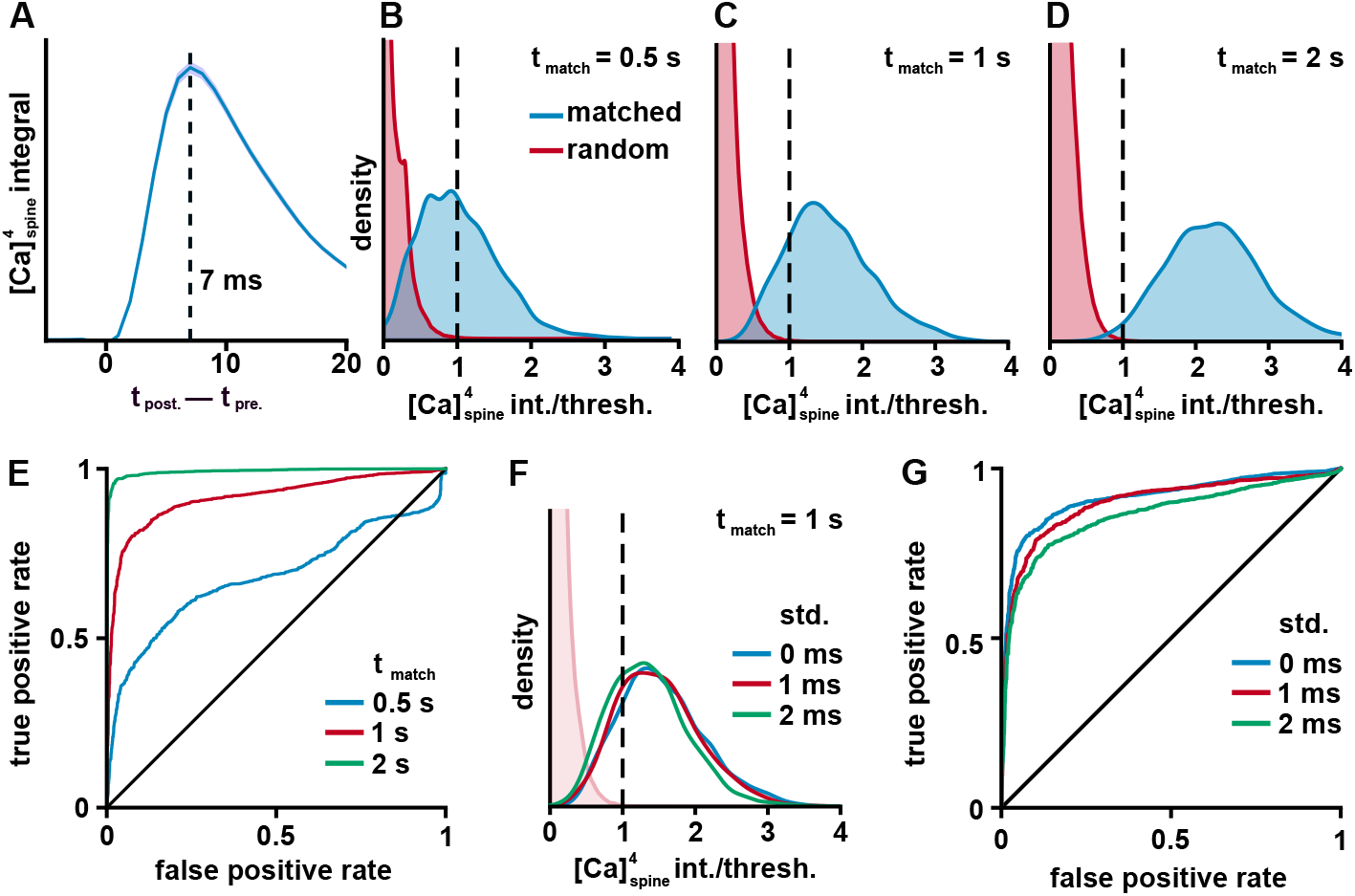
Testing if a threshold can distinguish between matched and random spike trains. **A)** The average over 300 simulations of the integral of the fourth power of the calcium concentration as a function of the time separation between pre- and postsynaptic spikes. Maximum at 7 ms. **B-D)** Kernel density estimate of the distribution of calcium integrals for matched spike trains (blue) and random spike trains (red). Match windows of size 0.5, 1, and 2 s shown. The calcium integrals were normalized by a threshold that rejected all non-matched spines 90% of the time (dotted line). **E)** Receiver Operator Characteristic (ROC) plot of the true positive rate vs. false-positive rate for various calcium integral thresholds. Note that the false-positive rate is for any of the spines receiving random input to have crossed the thresh-old. The 45-degree line, representing no discrim-ination ability, is shown for comparison. **F)** Kernel density estimate of the distribution of calcium integrals for matched spike trains (blue) and random spike trains (faint pink) when a small jitter is added to the times of the matched spike train. The time offsets were drawn from a gaussian with standard deviation of 0 (blue), 1 (red) and 2 (green) ms. **G)** A ROC plot as in panel E, but for different amounts of spike-time jitter and a 1 s match window.

For the match-and-control principle to work, it is necessary that a threshold can distinguish between a matching spike train and a random spike train. While this is easy to achieve when comparing a single matching train with a single random train, it becomes non-trivial when there are many random trains (our model has 149), none of which should potentiate. There is thus an unavoidable trade-off between rejecting as many random spike trains as possible and accepting as many matching spike trains as possible.

In this study, we selected an arbitrary rule that our thresh-old should reject all 149 non-matched spike trains 90% of the time and we normalize the calcium integrals by this threshold to make it easier to compare different conditions. Because of this, in every plot, 1 is the threshold for potentiation. We considered three different matching window sizes, 0.5, 1 and 2 seconds. To compare the ability of a threshold to distinguish between matched and random spines, we selected one group of spines to have a matched spike train input and the rest to be random spike trains, as described above. We ran 1500 simulations and computed a kernel density estimation of the distribution of calcium integrals, as shown in Figure 3 panels B, C, and D. Unsurprisingly, as the window size increases, more and more of the matched spines cross the threshold. This effect can also be seen in the receiver operator characteristic (ROC) plot (Figure 3E), where longer matching win-dows push the curve to the upper left. Finally, using a 1s window, we tested if adding temporal jitter to the presynaptic spike times would have an effect on the performance of our threshold. As can be seen in Figure 3 panels F and G, shifting each spike by a random number drawn from a Gaussian with standard deviation of 1 or 2 ms only had modest effects on distribution of calcium integrals, but, as expected, made the random and matching distributions have more overlap.

With our threshold that rejects the random spike trains 90% of the time, we found that the matched spines would cross threshold 45%, 82% or 98% of the time for 0.5, 1 or 2 second matching windows. Adding noise to spike trains in the 1 second window decreased the true positive rates to 78%, and 73% for 1 and 2 ms standard deviation noise, respectively. As should be expected, using longer windows and less noise allows for easier separation of the matched keys from random keys.

Note that, although we did not explore it in this study, increasing the number of axonal inputs to the dendrites likely increases the false positive rate. This can happen in two ways: First, with more random inputs, it becomes increasingly likely that a random spike train is very close to a matched spike train. Having similar keys can happen in an attention layer as well, but a major difference between a machine learning implementation and our system is that calcium is a noisy measure of spike train similarity, making it impossible to distinguish between nearly and exactly matching spike trains, even with a high threshold. Second, there is a small chance that non-matching spike trains will produce large amounts of calcium entry because of random fluctuations of the dendritic membrane or even spurious dendritic spikes. This issue has no analogue in transformers, but is thankfully only a rare occurrence in our model.

Because both issues have a roughly independent chance of happening for each axonal input, we expect that the true positive rate will decay exponentially to zero (holding the false positive rate constant) as the number of spines and presynaptic axons increases. We thus speculate that it would be difficult or impossible to implement our scheme, without major modifications, if the apical dendrite tuft received thousands of independent axonal inputs.

### The control phase

To potentiate matching spines, we multiplied the AMPA-component of the EPSP size by a sigmoid shown in Figure 4A, increasing it by a factor of eight when the calcium integral crossed our threshold. The baseline and potentiated EPSP sizes were tuned by hand to ensure that potentiated spines were able to drive dendritic spikes, while random fluctuations in the membrane potential rarely caused them. Because it is unrealistic for EPSP sizes to instantaneously increase, we added a time lag to the effects of the sigmoid with a time constant of 500 ms. This time constant was arbitrary, and we found that it had little effect on our simulations, though a minor benefit of the lag was that it reduced premature potentiation during the matching phase as well as spurious potentiation during the control phase when non-matched values happened to coincide with the value of the matched spine.

**Fig. 4.**
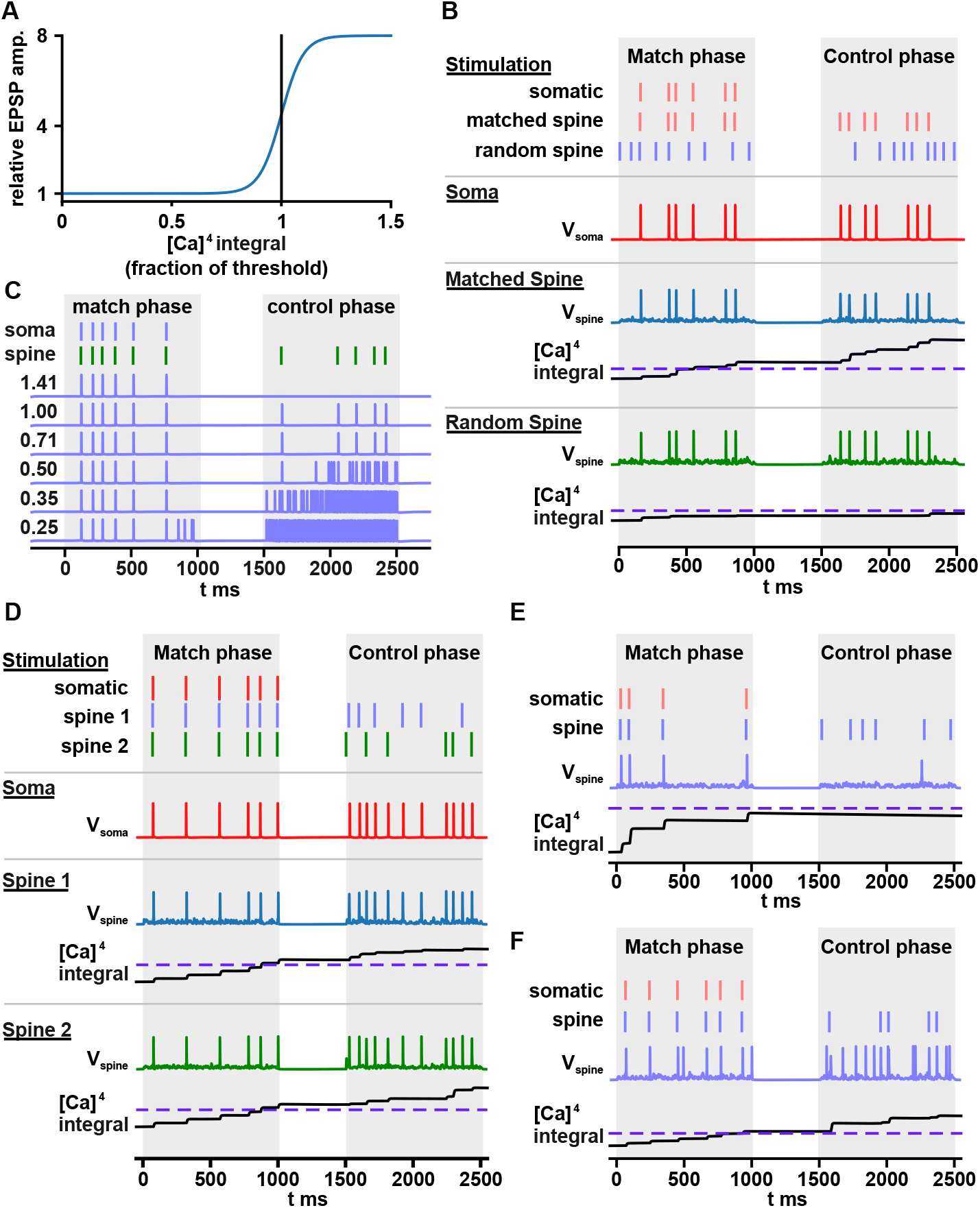
Simulations with potentiation and a control phase. **A)** Plot of the sigmoid used for potentiation of spine EPSPs as a function of integrated calcium. The threshold is shown as a vertical line. **B)** An example simulation showing a spine receiving input from an axon whose spikes coincided with the timing of BAPs (matched spine) and a second spine which received inputs from one of the other 149 axons with random spike trains (random spine). The random spine was selected as the most potentiated of all spines not receiving input from the matched axon. The somatic stimulation was only delivered during the match phase, but somatic spikes appear in the control phase that coincide with inputs to the matched spine (somatic voltage, red trace). The random spine did not elicit any somatic spikes. The integral of the fourth power of calcium is shown for both spines and the threshold is shown as a dotted blue line. Only the matched spine crossed the threshold. **C)** A set of repeated simulations with identical somatic and dendritic stimulation but with varying thresh-olds for potentiation. Only one of the 150 axons had a spike train that matched the BAPs, while the rest had random spike trains. Voltages are recorded in the soma. The numbers on the left represent fractions of our standard threshold that rejects all random axonal inputs 90% of the time. Threshold fractions are separated by factors of square root of 2. **D)** An example simulation where two spines had inputs from axons whose spike trains matched with the somatic stimulation. Both spines were able to transmit spikes to the soma producing an output that was the sum of the two input spike trains. **E)** Failure example 1: Despite one of the axons having a spike train that matches with the BAPs, the spines that receive this axonal input do not potentiate strongly enough to trigger dendritic spikes. **F)** Failure example 2: Multiple spines potentiated, only one of which had axonal input that matched the BAPs. Note that the matched spine was able to produce dendritic spikes, but most of the dendritic spikes originated from one of the other spines.

A typical simulation is shown in Figure 4B, which shows a spine receiving input from an axon whose spike train matches with the somatic spike train. After the match phase, no somatic stimulation is delivered, but the soma is driven to spike in time with the pre-dendritic axon’s action potentials, showing that the axon is able to control somatic activity. Also shown is the second-most-potentiated spine that received input from a pre-dendritic axon with a random spike train, but there is no correlation between this axon’s spike train and the somatic spike train. In our simulation, we left a small gap of 500 ms between the control and matching phases, but this is not necessary. We expect that in a real neuron, the control phase would begin immediately, but found that including this gap made it easier to understand our plots.

To see the effect of the calcium integral threshold, an identical set of pre- and postsynaptic inputs were simulated with different thresholds (Figure 4C). At the highest threshold, no potentiation occurs, while at lower thresholds spurious spikes occur during the control phase. At the lowest threshold, a cascading disaster occurs as more and more spines potentiate, and the dendrites are driven to spike wildly. In a more detailed ion-channel model, we expect that the dendrite would become persistently depolarized in this condition, rather than support ultra-high-frequency spiking. Regardless, we do not expect to see such behavior in real neurons as homeostatic processes in the neuron could detect excessive depolarization or spiking and increase the spine potentiation threshold to prevent it in the future.

We also examined the case where two groups of spines match with the BAPs pattern as shown in Figure 4D. In some such cases, as shown here, the spikes of the presynaptic spike trains are simply added together, but we note the chance of both trains crossing the threshold is the square of the chance of one of them crossing, making this occurence less likely than a successful potentiation of a single spine group. We note, though, that, when multiple spines are weakly potentiated, they will continue to potentiate during the control phase, since they are often depolarizing the spine enough to open NMDA channels. This continuing potentiation during the control phase is interesting as it translates the quality of the match into a non-trivial time-dependence, though we are unsure if it has biological relevance.

We also observe cases in which an axon with a spike train matched to the somatic activity fails to take control of the postsynaptic output. In Figure 4E, we see a failed potentiation of a spine that barely missed crossing the threshold (dotted line). In Figure 4F, we see an example where one of the random spines also potentiated. In this case, spikes originating from the spines receiving matched input only account for a small fraction of the spikes generated.

To evaluate the overall performance of the model, we performed 1000 simulations for each of the three match window sizes, 0.5, 1 and 2 s. In each simulation, one of the axonal inputs was again randomly selected to match with the somatic spike pattern during the matching phase, while the other 149 axons were drawn from a 6 Hz Poisson process. We then characterized each of the matched axon’s presynapatic inputs during the control phase as “successful” only if it was followed within 20 ms by a somatic spike. In contrast, each somatic spike that was not preceded by an input from the matched axon in this same window was considered “spurious”. The percent of successful and spurious somatic spikes is shown in Figure 5A.

**Fig. 5.**
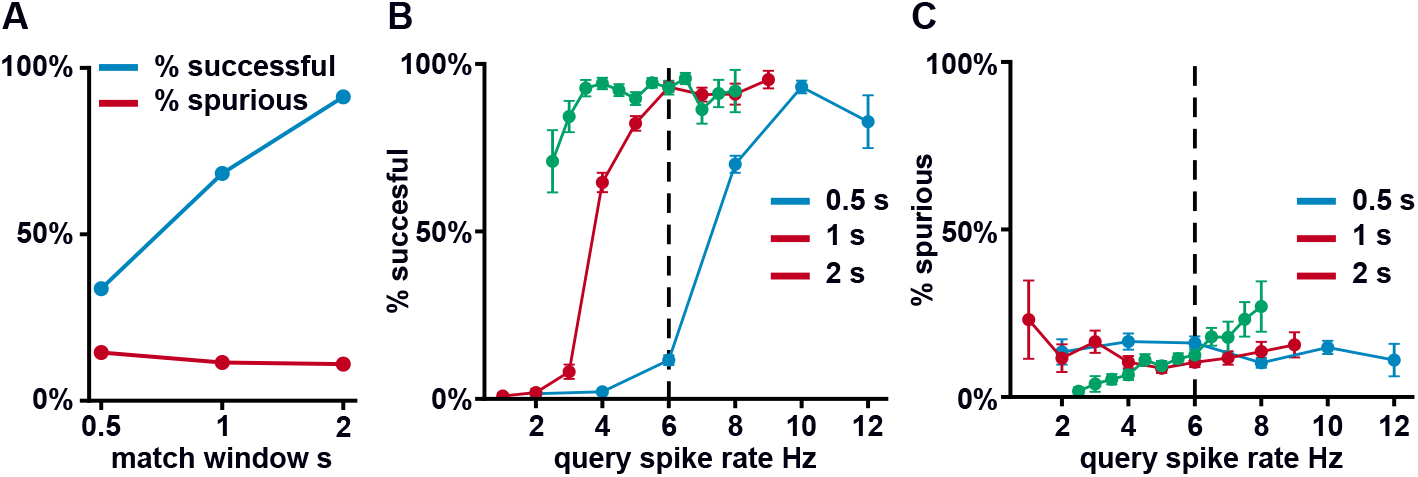
Performance of the model during the control phase. **A)** The results from 1000 simulations of the model for each of the three window sizes are shown. In each simulation, one axonal input was matched with the somatic spiking pattern and the rest were random. The percent of axonal inputs from the matched axon that produced a somatic spike is shown in blue (% successful). The percent of spikes that could not be explained by axonal stimulation from the matched axon is shown in red (% spurious). **B&C)** The percent of spikes that were successful and spurious is broken down by query spike rate.

We observed that the spike rate of the query had a strong influence on the percent of successful spikes, especially for the 0.5 and 1 s match windows (Figure 5B), while having a more modest effect on the percent of spurious spikes (Figure 5C). This finding is not surprising, since each somatic spike gives a chance for NMDA receptors to open and allow calcium entry. When there are fewer spikes, less calcium will enter each spine, decreasing the chance that the spine will cross the threshold for potentiation. It is thus very natural to fix the spike rate of the query to be the average spike rate of the axonal inputs. Indeed, doing so for 1 s match windows raises the percent of successful spikes up to that of the 2 s windows.

## Discussion

The match-and-control principle is a simple mechanism for implementing something akin to the attention layers of transformers. Attention is computationally expensive, but our model allows for large numbers of key and query comparisons because it uses one of the smallest computational units of the brain, small groups of spines, to perform each overlap. Here we briefly discuss the limitations of this proposal as well as possible circuits in the brain where it might plausibly occur.

In our view, the most troublesome aspect of our proposal is the large potentiation of synapses required for the axonal inputs to control somatic activity. In our model we used a factor of eight increase in EPSP size, which is much larger than what is typically seen following LTP induction. Remarkably, such large increases in EPSP size have been observed in short-term potentiation as far back as Bliss and Lomo (39), but this effect is believed to be presynaptic (40) and thus cannot depend on the synchrony of pre- and postsynaptic spikes. One possible mechanism that we have contemplated is that presynaptic strengthening occurs broadly across the dendritic tree following the match phase, allowing presynaptic spikes to drive dendritic spiking, but that postsynaptic depression of non-matching spines occurs simultaneously, blocking this effect for all but a few spines. A similar idea is that the spiking patterns of the value could differ qualitatively from those of the key. If the value spike train consists of bursts of action potentials, this would make it easier for single axons to initiate dendritic spikes, but would again require depression of non-matching synapses.

Calcium-dependent ion channels offer another mechanism for depolarizing the spine. Notably, transient receptor potential (TRP) channels such as TRPM4 (41) and TRPM5 (42, 43) that are impermeable to calcium, could allow for rapid depolarization of dendrites following NMDA-dependent calcium entry, allowing for easier dendritic spike initiation. Calcium-dependent potassium channels, such as SK channels, are found in spines (44), and could depress the voltage in spines with low levels of calcium entry. Considerable care would have to be taken in the use of these channels as any persistent increase in membrane potential could cause NMDA channels to open in the absence of BAPs. Nonetheless, we believe that careful use of them could reduce the size of the potentiation needed for our simple model.

A related issue is the ability of dendritic spikes to faithfully propagate to the soma. We found it necessary to carefully design the interface between the small dendrites of the tuft and the much larger apical dendrite to prevent dendritic spikes from dying out. We resolved this problem by adding a region of somewhat higher sodium channel density. In large pyramidal neurons this active zone might be replaced by region of high-voltage-activated calcium channels, such as L-channels, but the slower kinetics of such channels may have difficulty in faithfully transmitting the individual spikes in a spike train.

This may make it advantageous to implement our mechanism in pyramidal neurons with narrower apical dendrites than we used in our model or to use a value consisting of bursts of spikes that may be easier to transmit to the soma.

While these problems may appear as great challenges for our design, we note that Larkum, in his apical dendrite “manifesto” (45), suggested that small clusters of spines on layer 2/3 neuron apical dendrites might “with explosive impact dictate the firing of the neuron”. In his formulation, this effect arises because broad dendritic inhibition silences most of the spines, while sparing a small number of them that are depolarized by NMDA spikes. We extend this hypothesis by observing that, if some Hebbian mechanism can be discovered that selects which groups of spines are allowed to control the neuron, it would allow for an attention-like computation in cortex.

Another major objection that one might raise to our proposal is that NMDA-dependent plasticity already has a purpose in the brain, LTP, and our new mechanism would seem to crowd in on previous theories of learning. However, transient increases in synaptic strength that are larger than what will remain asymptotically are common in LTP experiments and many theories have been proposed that might use them (23, 40). Furthermore, if the synapses in our model undergo LTP slowly after each match phase, this would imply that axons that commonly match with the activity of a neuron become preferred in future comparisons. While this would break the democratic matching process of attention layers in transformers, such a bias could be advantageous. Since our matching procedure involves a noisy comparison between spike trains (because of random membrane fluctuations) comparing thousands of keys with a query may be nearly impossible in vivo and a mechanism, like LTP, that selects likely candidates for matching would be beneficial. A far more challenging question is how a neural implementation of transformers could be trained through experience. This problem remains unsolved for traditional models of neural circuits as well, but we note that there are promising methods for training artificial spiking transformers (18), even if they have not yet achieved the same performance as standard rate-coded models.

An additional puzzle is what mechanism might synchronize the arrival of the keys with the start of the queries. While it is tempting to assume that cortical rhythms might play a role, we have found that match windows around 0.5 - 2 s are necessary for reducing noise in the spike train comparisons, a timescale much longer than the cycles of theta, beta or gamma rhythms found in the cortex. Bouts of cortical activity lasting around a second are observed following sensory stimulation, or during the “UP states” that occur in both sleeping and awake but resting animals (46, 47). The match phase may be triggered off the start of these periods of high activity.

Finally, we turn to the question of which neural populations in the brain might implement our program. We selected a pyramidal neuron for our model because the apical dendrite is a natural mechanism for transmitting dendritic spikes to the soma. However, large layer V pyramidal neurons in particular have several disadvantages. First, they are known to have dendritic spikes that do not reach the soma and their dendrites are controlled by complex dendritic inhibition (48, 49). Second, BAPs have been observed to broaden temporally as they reach the distal tuft, likely due to calcium channel participation, which would harm our precise temporal matching (49). Finally, they are the output neurons of cortex, and, as was discussed above, in many transformer models, attention layers are typically only highly selective in the middle layers of transformers, not in their input or output layers.

We thus conjecture that if our mechanism occurs, it is more likely to be found in either thin tufted layer V neurons (50, 51) or in layer 2/3 neurons both of which project primarily within the cortex. Layer 2/3 neurons, in particular, may have the best ability to propagate BAPs to the dendrites for matching, and dendritic spikes to the soma for transmission, due to their relatively short apical length (52).

## Methods

All computations were performed on a desktop PC with a 12 core Intel Xeon Processor using the Python interface for the NEURON simulator. All python code and NEURON channel MOD files, including code for generating the data and figures are available from a public GitHub repository, https://github.com/iellwood/MatchAndControlPaper. The repository includes instructions for running the code and a script that prints the NEURON model’s parameters, including the conductances of the individual ion channels.

## ACKNOWLEDGEMENTS

We would like to thank Andrew Bass, Ronanld Hoy, Christiane Linster, Monzilur Rahman, Weinan Sun, Melissa Warden and Ronald Harris-Warrick for discussions and comments on the manuscript. This research was supported in part by a BBRF Young Investigator Grant.

## AUTHOR CONTRIBUTIONS

E. performed all work in the paper, including simulations, analysis and writing of the manuscript.

